# Shotgun metagenomics unravels higher antibiotic resistome profile in Bangladeshi gut microbiome

**DOI:** 10.1101/2023.03.02.530749

**Authors:** Arittra Bhattacharjee, Tabassum Binte Jamal, Ishtiaque Ahammad, Zeshan Mahmud Chowdhury, Anisur Rahman, Gourab Dewan, Shiny Talukder, Mohammad Uzzal Hossain, Keshob Chandra Das, Chaman Ara Keya, Md Salimullah

**Affiliations:** Bioinformatics Division, National Institute of Biotechnology, Ganakbari, Ashulia, Savar, Dhaka, 1349, Bangladesh; Department of Mathematics and Natural Sciences, BRAC University, 66 Mohakhali, Dhaka 1212, Bangladesh; Rangamati Medical College, Hospital Road, Rangamati-4500, Bangladesh; Molecular Biotechnology Division, National Institute of Biotechnology, Ganakbari, Ashulia, Savar, Dhaka, 1349, Bangladesh; Department of Biochemistry and Microbiology, North South University, Bashundhara, Dhaka, 1229, Bangladesh

**Keywords:** Antibiotic resistance, resistome, shotgun metagenomics, mobilome, LMICs

## Abstract

Antibiotic resistance management is a challenging task in Low and Middle-Income Countries (LMICs) such as Bangladesh. Improper regulation and uncontrolled spreading of Antibiotic Resistant Genes (ARGs) from LIMCs pose a great threat to global public health. The human gut microbiome is a massive reservoir of Antibiotic Resistant Genes (ARGs). In this study, we unraveled the ARGs in the gut microbiome of the Bangladeshi population and compared them with several other countries around the world. Here, 31 fecal samples from different ethnic groups living in Bangladesh namely Bengali (n=9), Chakma (n=6), Khyang (n=5), Marma (n=6), and Tripura (n=5) were collected. Shotgun metagenomic sequencing method was implemented for revealing the ARGs. The resistome profiling was executed on three levels-the total microbiome, the plasmidome, and the virome. In all three levels, samples from Bangladeshi cohorts showed higher ARG profiles compared to foreign samples. On average, the number of ARGs in the Bangladeshi samples ranged between 75.11 and 88. Among them, class C beta-lactamases, quinolone resistance genes, and tetracycline efflux pumps were relatively more abundant. Additionally, the MexPQ-OpmE drug resistance pathway was found to be more prevalent. Findings from our study suggest that the spread of antibiotic resistance within the Bangladeshi population is being facilitated by the gut microbiome especially via the mobilome. Therefore, strict regulation on antibiotic usage is necessary to halt the spread of ARGs.

## Introduction

Antibiotics are the most potent life-saving drugs developed in the twentieth century, saving millions of lives from bacterial infections and reducing significant public health concerns such as infectious diseases, maternal and infant mortality (1). Based on worldwide pharmaceutical sales statistics in 76 countries, antibiotic usage in defined daily doses increased by 65 percent from 2000 to 2015, with predicted consumption to increase by more than 200 percent by 2030 (2). Antibiotic treatment was formerly more common among high-income countries; but, due to an increase in revenue, antibiotic consumption increased dramatically in low and middle-income countries including India, China, Pakistan, and Bangladesh (3,4). Despite the fact that antibiotics are a new addition to humanity’s overall history, the capability of bacteria to resist various antibiotics is ancient (5). (Peterson & Kaur, 2018). Antibiotic resistance in pathogenic bacteria has appeared as a major public health concern in recent decades addressing economic loss, treatment failure, and challenges in global health security (5). In the United States, about 35,000 people die each year from antibiotic-resistant diseases (6).

Antibiotics are classified into two groups based on their functions: bactericidal antibiotics kill bacterial cells while bacteriostatic antibiotics inhibit the growth of microbial cells. Some of the broadly used bacteriostatic antibiotic groups are Tetracyclines (e.g. doxycycline, minocycline) whereas common bactericidal antibiotics are Aminoglycosides (e.g.gentamicin, neomycin), Beta-lactams (e.g. penicillins, carbapenems, cephalosporins) or Fluoroquinolones (e.g. Ciprofloxacin) (7). The continuous growth of microorganisms even in the presence of antimicrobial agents has been one of the most important and persistent driving forces for antibiotic discovery. The potency of the bacteria to change its physical appearance and genetic makeup has led to resistance to almost all available antibiotics in clinical practice. Gram-negative bacteria can utilize their outer cell membrane to block the antibiotic entrance, while other microorganisms employ efflux pumps, manufacture various proteins and modify their targets to get rid of the opponent molecule (8). In the process, they obtain antibiotic resistance genes (ARGs) that encode intrinsic antibiotic resistance mechanisms. Resistance is considered a natural phenomenon of bacteria; however, the overuse/misuse of antibiotics, as well as the slow pace at which new medicines are discovered has expedited the dissemination of the resistance gene. ARGs can be shared among bacterial organisms via Horizontal Gene Transfer (HGT) through conjugation or transduction commencing rapid development of multi-drug resistance in bacteria. Plasmid, transposons, and mobile integrons act as vital mediums in gene transfer. Studies on lateral gene transmission demonstrated gene transfer to be 25 times more in human-associated bacteria than in other bacteria in the environment (9).

Antimicrobial resistance is the greatest threat to Southeast Asian countries, according to the World Health Organization (WHO) (10). Almost all Asian countries, including Sri Lanka, China, South Korea, Japan, Hong Kong, Taiwan, Thailand, Singapore, and India, harbor methicillin and erythromycin-resistant bacteria. The dissemination of antibiotic-resistant bacteria from Asia to other nations is a regular occurrence. New Delhi metallo-beta-lactamase-1 (NDM-1) generated in India spread to the United Kingdom, Sweden, Austria, Belgium, France, the Netherlands, Germany, the United States, Canada, Japan, China, Malaysia, Australia, and Korea (11). Colistin-resistant Enterobacteriaceae originated in China in 2016 and has subsequently spread to more than 30 nations (12). Bangladesh being in close proximity to India is also susceptible to antimicrobial resistance. Besides, proper maintenance of hygiene cannot be assured all the time in this country, and therefore, antibiotics are an important resource to control different epidemics, nosocomial infections, and post-surgery sepsis (13). Concurrently, the availability of antimicrobial medicine as an over-the-counter drug with influential unethical pharmaceutical practices and irresponsible antibiotic usage in agriculture and farming has escalated its exploitation in the nation (4). A two-fold increase in multidrug resistance has been observed between 2015 and 2019, amid which *E. coli* and *Staphylococcus aureus* were found to be the most prevalent multidrug-resistant pathogens (14). In another study, typhoid patients were found to be resistant to second-line therapy (ciprofloxacin) conducted in the Chittagong division in 2003 (15). The efficacy of various antibiotics like ampicillin, sulfamethoxazole-trimethoprim, and tetracycline no longer show efficacy as before. Chittagong division encompasses three districts of Chittagong Hill Tracts, considered the only extensive hill area in Bangladesh. It consists of 1.6 million people, of which about 50% belong to small ethnic groups (16). The prevalence of antimicrobial resistance in tribal people has been largely ignored. A recent study stated that these indigenous groups self-prescribe various antibiotics which is boosting the increasing rate of antibiotic resistance (17).

The human gut is a reservoir of interdependent microorganisms that participates in digestion, immune system maintenance, metabolic reactions, and protection from pathogens (18). Several studies showed that antibiotics can disrupt the taxonomic compositions of the human microbiota. The gut viral components are highly dominated by the bacteriophages or phages known to have significant roles in shaping microbial composition (19). Polymerase chain reaction on viral DNA showed ARGs to be resistant to various antibiotics including lactamase, quinolone, and vancomycin. ARGs were found to be present in the human gut for several years even after short-term antibiotic intake (20). In this study, we explored the resistome profile of different indigenous groups of Bangladesh and the mainstream Bengali population using shotgun metagenomics. Afterward, we compared them with the healthy population from 11 countries across the world. The resistome pattern was similar among all the Bangladeshi but significantly distinct from the other cohorts.

## Methodology

### Sample Collection

A total of 31 fecal samples were collected from four ethnic groups (all applicable international, national, and/or institutional guidelines regarding human samples were followed; ethical approval was given by the Ethical review board of the National Institute of Biotechnology) including Chakma, Marma, Khyang, and Tripura from Rangamati, Chittagong hill tracts (22.6533° N, 92.1789° E) and Bengali from Dhaka (23.9536° N, 90.1495° E), Bangladesh. Participants did not have any recent records of antibiotic consumption. The samples were transported in an ice box from the collection site to the National Institute of Biotechnology and immediately stored at -80°C before DNA extraction.

### Data Collection from other studies

To compare our samples with other studies around the world, we obtained raw metagenomic reads from the National Center for Biotechnology Information (NCBI) Sequence Read Archive (SRA) that were submitted after sequencing. Samples/reads were collected from Australia (n=8), Cameroon (n=10), China (n=10), Denmark (n=9), Germany (n=10), Hong Kong (n=9), India (n=10), Italy (n=5) and Korea (n=10), their NCBI Bioproject numbers are PRJNA503909, PRJEB27005, PRJNA624763, PRJEB2054, PRJEB48605, PRJEB48269, PRJNA39711, PRJNA386422, PRJNA743718, PRJNA48479 respectively. Moreover, reads were obtained from NIH Human Microbiome Project (HMP) (https://hmpdacc.org/). Six healthy samples (from the USA) were collected from the HMP1 portal.

### DNA extraction and metagenomics Sequencing

DNA was extracted from fecal samples using the Invitrogen™ PureLink™ Microbiome DNA Purification Kit. Here, 0.2±0.05 g stool samples were transferred to special beads containing tubes and went through PureLink™ spin-column technology. DNA concentration and purity were measured using Thermo Scientific™ NanoDrop™ 2000/2000c and stored at -20°C. For metagenomics sequencing, library preparation was done using TruSeq® DNA Library Prep Kits. Afterward, sequencing was performed in Illumina HiSeq 2500.

### Raw sequence processing and assembly

The Fastq raw reads were aligned with Burrows-Wheeler Aligner (BWA) against *Homo sapiens* (assembly GRCh38.p13) to remove human DNA contaminants (21). The unaligned reads were trimmed using Trimmomatic (Bolger et al., 2014). The trimmed reads underwent *de novo* assembly using MEGAHIT (22).

### Virome identification

To sort out the viral sequences, the assembled contigs were screened using DeepVirFinder (https://github.com/jessieren/DeepVirFinder). DeepVirFinder implemented a reference and alignment-free machine learning method to differentiate the viral sequences in the metagenomic contigs via convolutional neural networks (CNN) based deep learning (23). DeepVirFinder gave a score to every contig. Contigs with ≥ 0.9 score and p-value with ≤ 0.05 were considered viral contigs.

### Classification of plasmidome

The plasmidal contigs were classified using PlasClass (https://github.com/Shamir-Lab/PlasClass). PlasClass can classify short plasmidal sequences from metagenomic samples with improved and more reliable methods. Contigs with ≥ 0.9 score were selected as plasmidal contigs.

### Antibiotic resistome detection

The resistant genes were identified from viral, plasmidal, and total contigs via ABRicate 0.5 (https://github.com/tseemann/abricate). This tool can use databases such as NCBI, CARD, ARG-ANNOT, Resfinder, MEGARES, EcOH, PlasmidFinder, Ecoli_VF, and VFDB. Here, the Resfinder database was implemented to find out the antibiotic resistance genes. Afterward, the contigs were translated using Prodigal (24). The gene count data from ABRicate went under log 10 transformation. The transformed datasets were implemented for Kruskal Wallis test followed by Pairwise Wilcoxon test and Spearman’s rank correlation. R version 4.2.2 (https://www.R-project.org) and ggplot2 package (https://ggplot2.tidyverse.org) were utilized for all statistical analyses and visualization (25). The antibiotic-resistant causing proteins were determined by fARGene (26). fARGene applies optimized gene models hence it can identify previously uncharacterized resistance genes with higher accuracy, even if their sequence similarity is low to known ARGs.

### Pathway analysis

To identify the drug resistance-related pathways, Prodigal annotated proteins were classified using Micorbeannotator. MicrobeAnnotator is a comprehensive functional annotation for microbial genomes that merges results utilizing KEGG Orthology (KO), Enzyme Commission (E.C.), Gene Ontology (GO), Pfam, and InterPro (27). From all the pathway tables, drug-resistant pathways were sorted out. Linear discriminant analysis (LDA) of the metabolic pathways was conducted using Linear discriminant analysis Effect Size (LEfSe) (28).

## Results

### Prevalence of ARGs was higher in the samples of Bangladesh

Antibiotic resistant gene (ARG) homologs were detected by ABRicate. ARGs were present in every Bangladeshi sample. Among all the local populations, the Khyang population had the highest number of ARGs identified. On average, 88 ARG were found in this population, with a standard deviation of 25.06. (**Table 1: SK1-SK17**).

**Table 1:** Bangladeshi ethnic cohorts and the distribution of antibiotic-resistant gene homologs in their total microbiome, plasmidome, and viraome as detected by ABricate.

| Sample No. | Country | Gender | Antibiotic Resistant Genes | Total Plasmidal Antibiotic Resistant Genes | Viral Antibiotic Resistant Genes |
| --- | --- | --- | --- | --- | --- |
| SB4 | Bangladesh(Bengali) | Female | 97 | 14 | 2 |
| SB7 | Bangladesh(Bengali) | Female | 72 | 11 | 5 |
| SB8 | Bangladesh(Bengali) | Female | 84 | 7 | 4 |
| SB1<br>4 | Bangladesh(Bengali) | Female | 72 | 9 | 0 |
| SB1<br>5 | Bangladesh(Bengali) | Male | 74 | 7 | 2 |
| SB2<br>1 | Bangladesh(Bengali) | Male | 66 | 7 | 2 |
| SB2<br>4 | Bangladesh(Bengali) | Male | 82 | 9 | 4 |
| SB2<br>5 | Bangladesh(Bengali) | Male | 63 | 8 | 0 |
| SB3<br>3 | Bangladesh(Bengali) | Male | 66 | 8 | 2 |
| SC2 | Bangladesh(Chakma) | Female | 103 | 17 | 1 |
| SC3 | Bangladesh(Chakma) | Female | 70 | 7 | 2 |
| SC4 | Bangladesh(Chakma) | Male | 71 | 9 | 4 |
| SC5 | Bangladesh(Chakma) | Male | 86 | 4 | 4 |
| SC6 | Bangladesh(Chakma) | Female | 72 | 5 | 5 |
| SC2<br>5 | Bangladesh(Chakma) | Female | 89 | 8 | 2 |
| SK<br>1 | Bangladesh(Khyang) | Male | 126 | 20 | 2 |
| SK<br>2 | Bangladesh(Khyang) | Female | 62 | 11 | 1 |
| SK<br>3 | Bangladesh(Khyang) | Female | 70 | 11 | 6 |
| SK<br>4 | Bangladesh(Khyang) | Male | 86 | 13 | 1 |
| SK<br>17 | Bangladesh(Khyang) | Male | 96 | 5 | 3 |
| SM<br>1 | Bangladesh(Marma) | Femal<br>e | 107 | 16 | 1 |
| SM<br>2 | Bangladesh(Marma) | Femal<br>e | 79 | 8 | 4 |
| SM<br>3 | Bangladesh(Marma) | Male | 80 | 9 | 1 |
| SM<br>4 | Bangladesh(Marma) | Femal<br>e | 94 | 8 | 2 |
| SM<br>5 | Bangladesh(Marma) | Male | 92 | 13 | 6 |
| SM<br>6 | Bangladesh(Marma) | Femal<br>e | 71 | 12 | 4 |
| ST1 | Bangladesh(Tripura) | Male | 83 | 7 | 8 |
| ST3 | Bangladesh(Tripura) | Femal<br>e | 66 | 4 | 2 |
| ST4 | Bangladesh(Tripura) | Female | 98 | 20 | 3 |
| ST6 | Bangladesh(Tripura) | Female | 93 | 9 | 7 |
| ST1<br>5 | Bangladesh(Tripura) | Female | 62 | 8 | 2 |

The second and third highest number of ARGs were found in Marma (87.17± SD 12.98) and Chakma (81.83± SD 13.2) respectively. Tripura samples showed 80.4± SD 15.98 ARGs and Bengali had 75.11 ± SD 10.83 ARGs. In Bangladeshi samples, Bengali groups had the lowest ARGs. The heatmap analysis (**Supplementary Figure 1**) demonstrated clusters where tet(W), tet(Q), tet(32), tet(37), catB4, VanR_G, tet(O), oqxA, tet(44), blaOXA-347, erm(X), aadA1, and many other ARGs were highly abundant in Bangladeshi population whereas lower abundance was observed in other population (**Supplementary Figure 1**). According to the Kruskal-Wallis test, the local and global groups have different distributions of ARG abundance. In general, Bangladeshi samples possessed higher abundance of ARGs compared to foreign samples (**Figure 3a**)(**Table 2**).

**Table 2:**
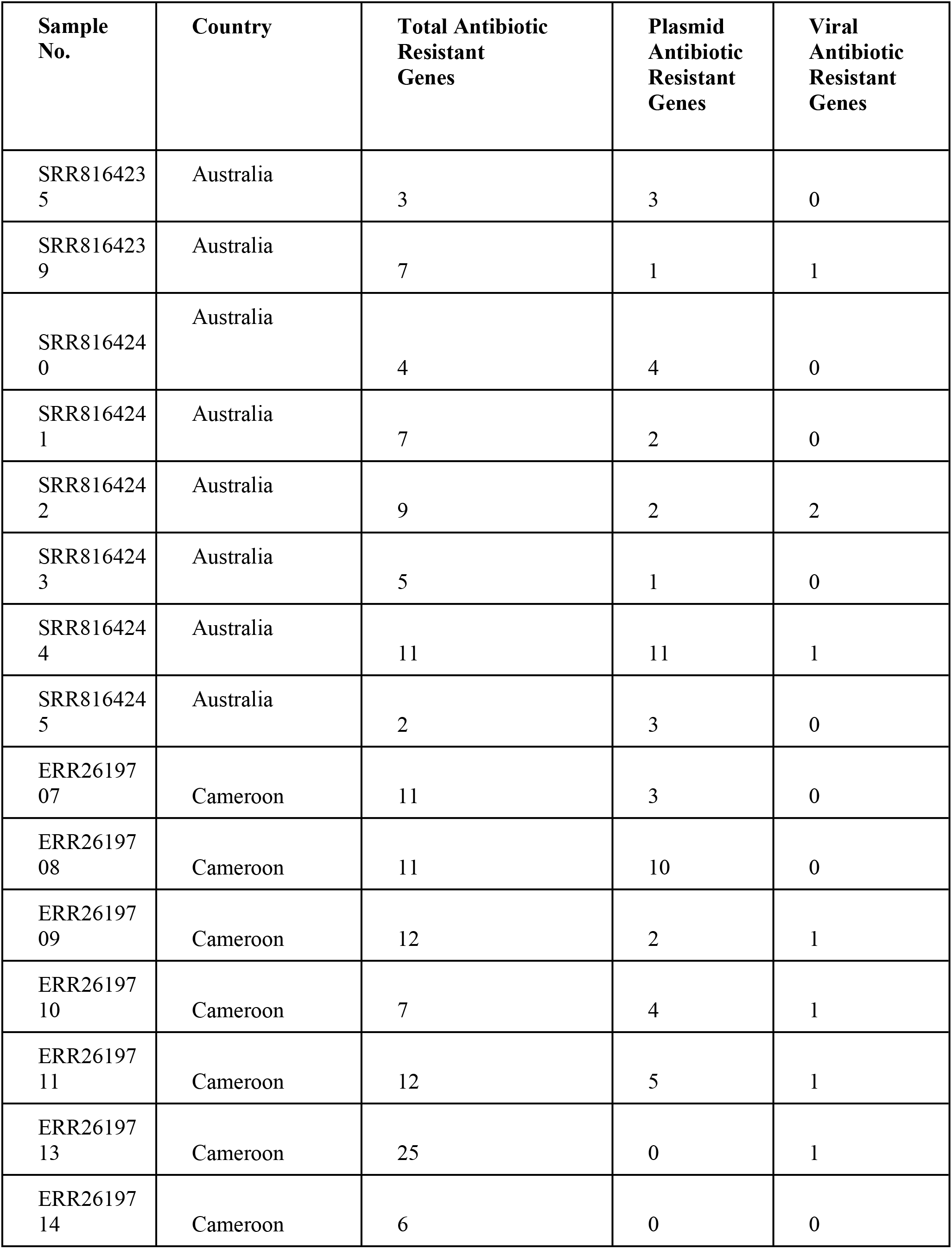
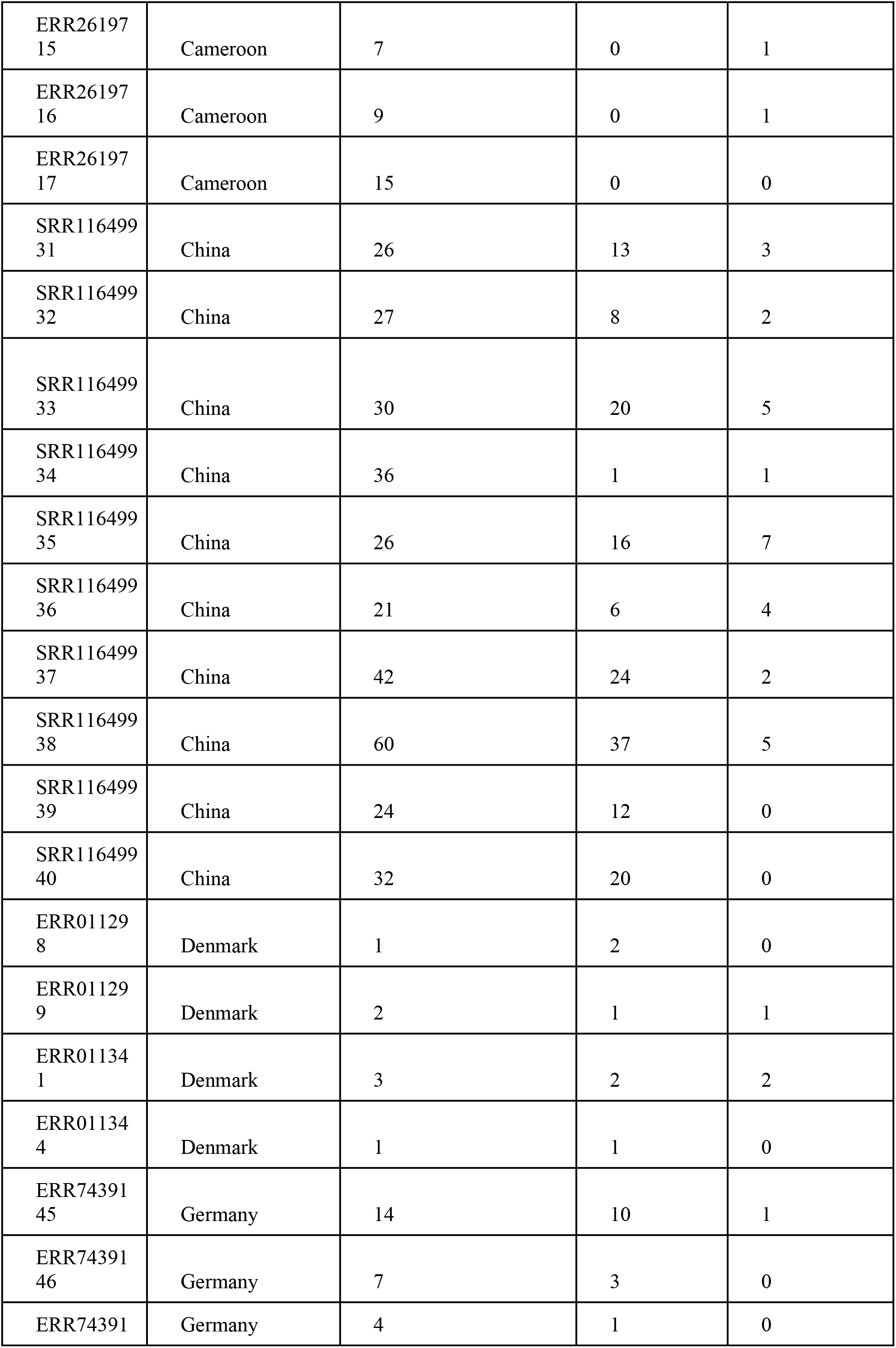
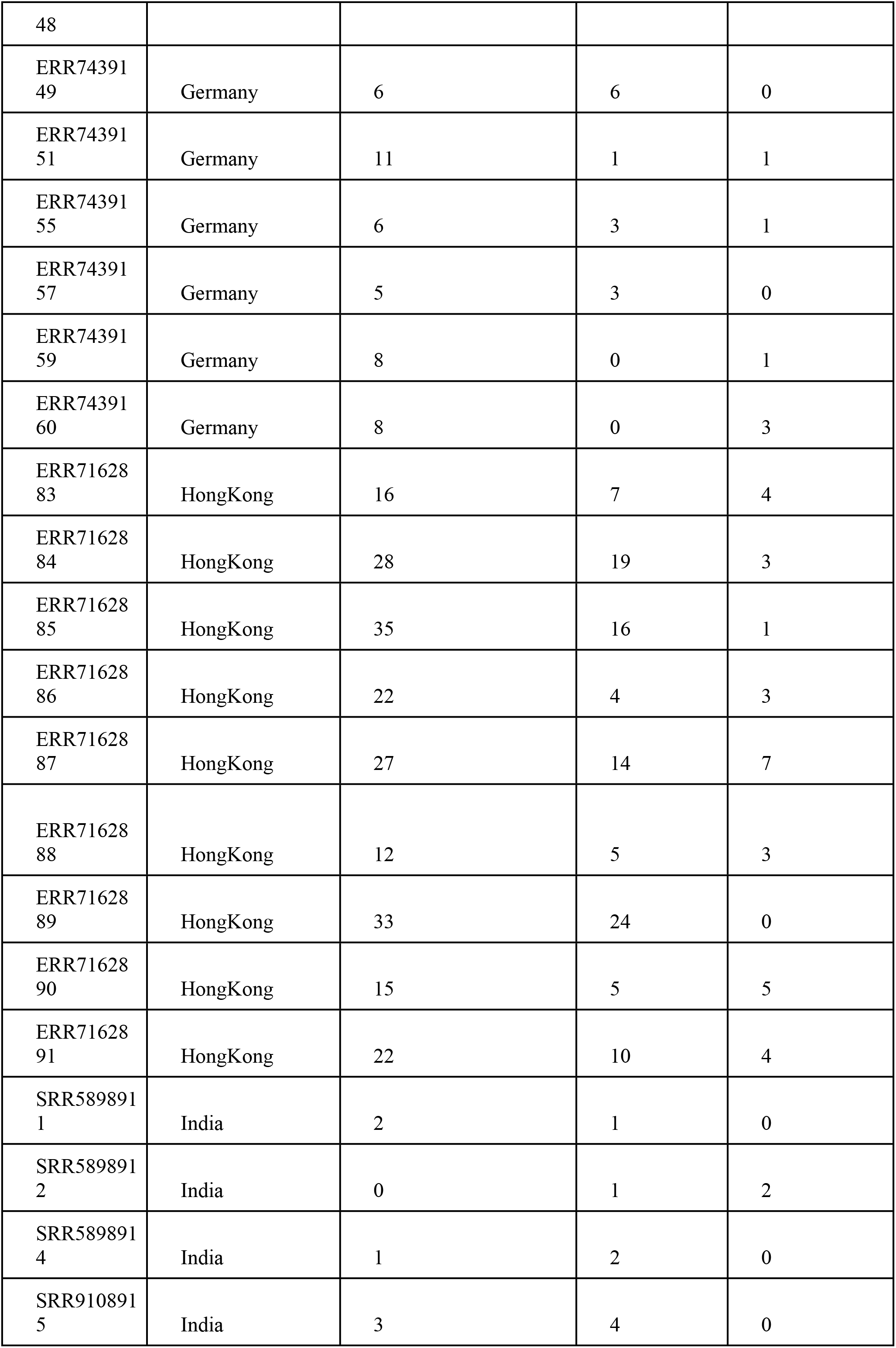
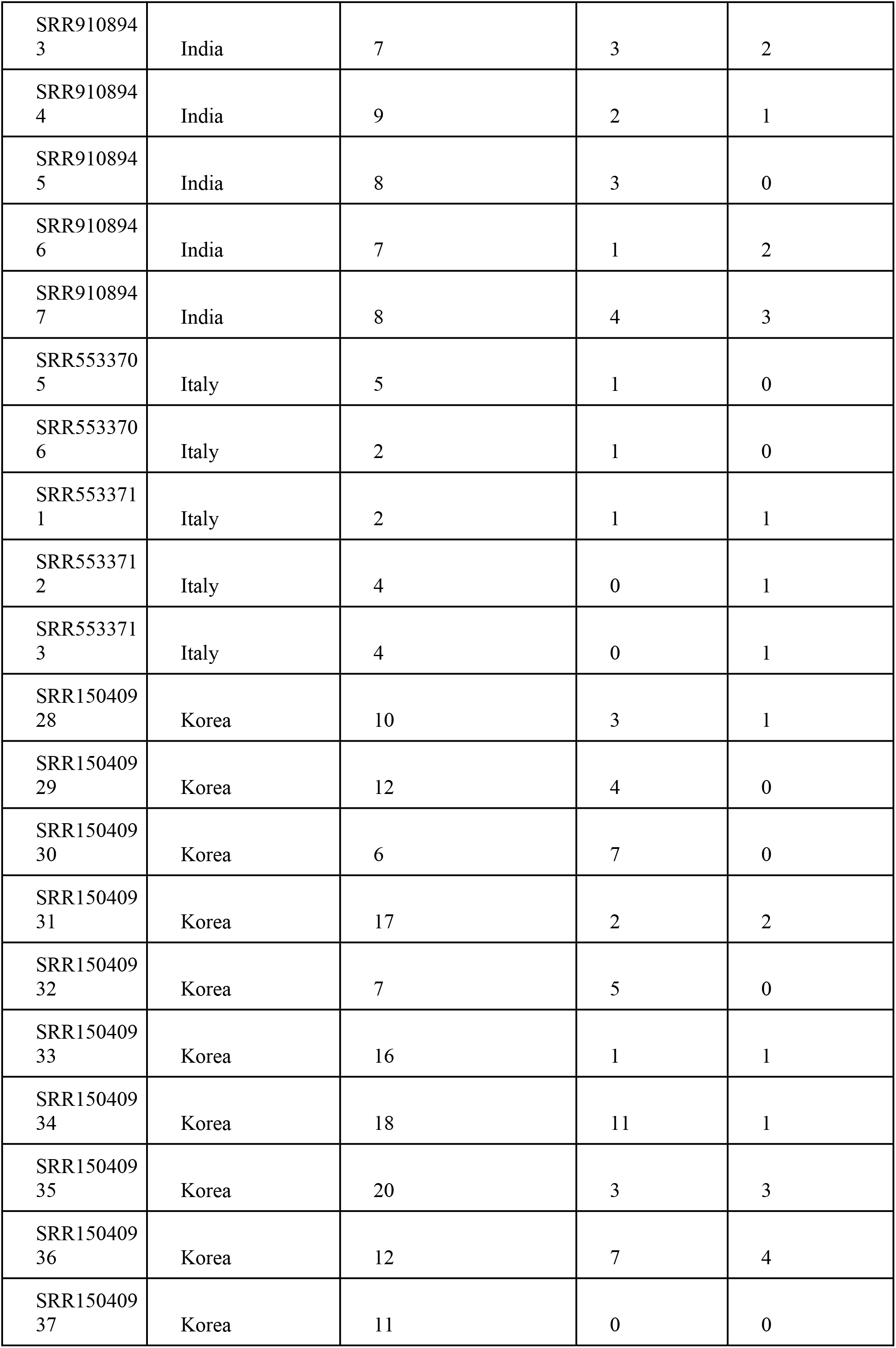
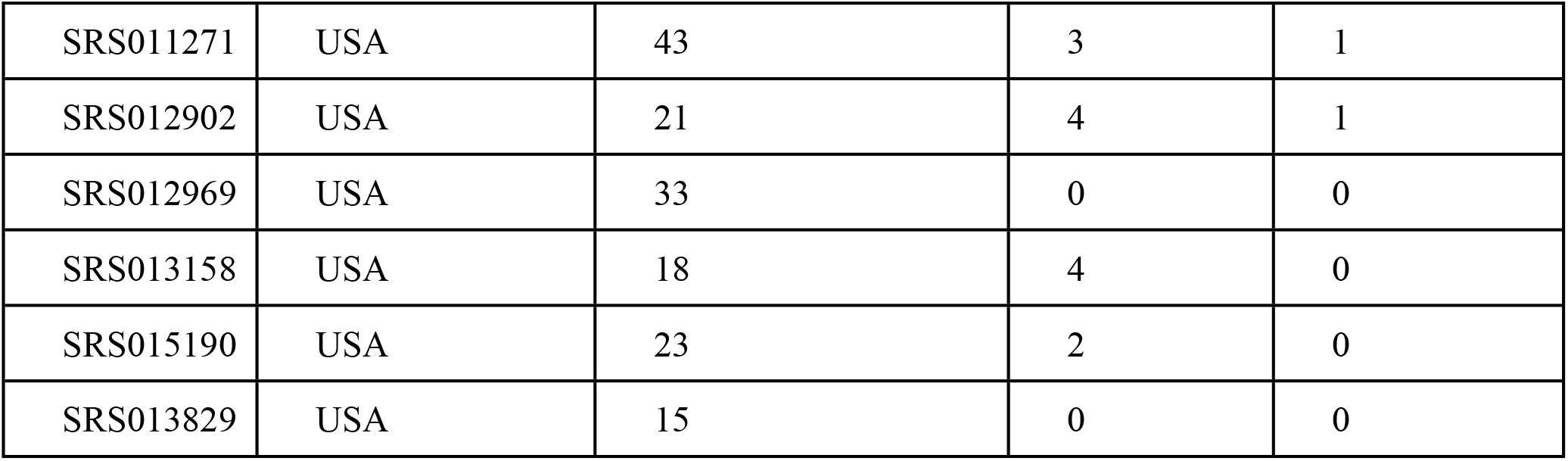
International cohorts and the distribution of antibiotic-resistant gene homologs in their total microbiome, plasmidome, and virome as detected by ABricate.

However, samples from Bangladesh are not statistically different (p-value> 0.5) (**Figure 3b**). The protein level identification from fARGene analysis showed Class_a and class_tet_rpg were highly prevalent in the Bangladeshi population. Interestingly, Class_b_1_2 was less abundant in the Bangladeshi groups compared to other populations and class_d_1 was absent (**Figure 2**).

**Figure 1:**
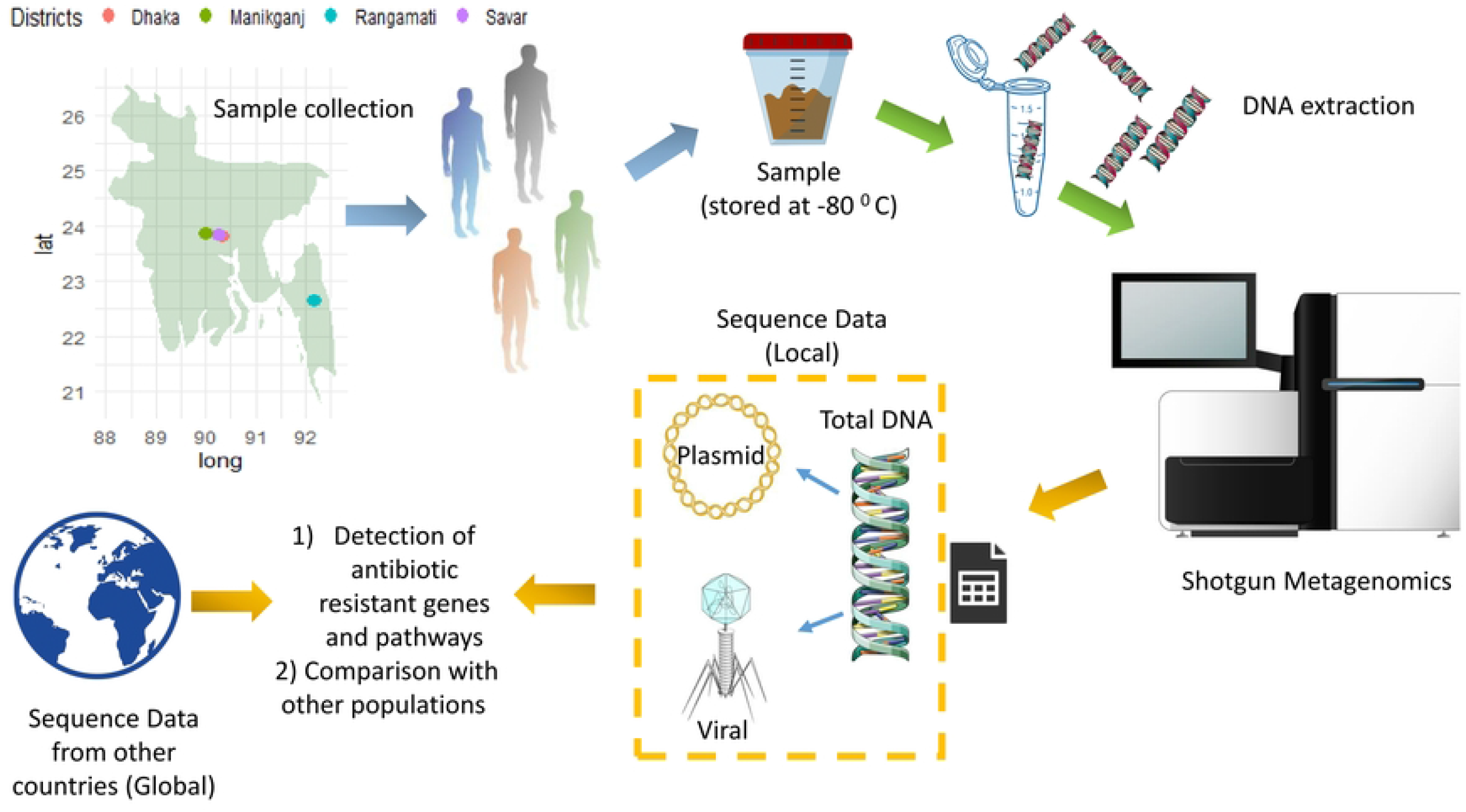
Schematic representation of the whole study.

**Figure 2:**
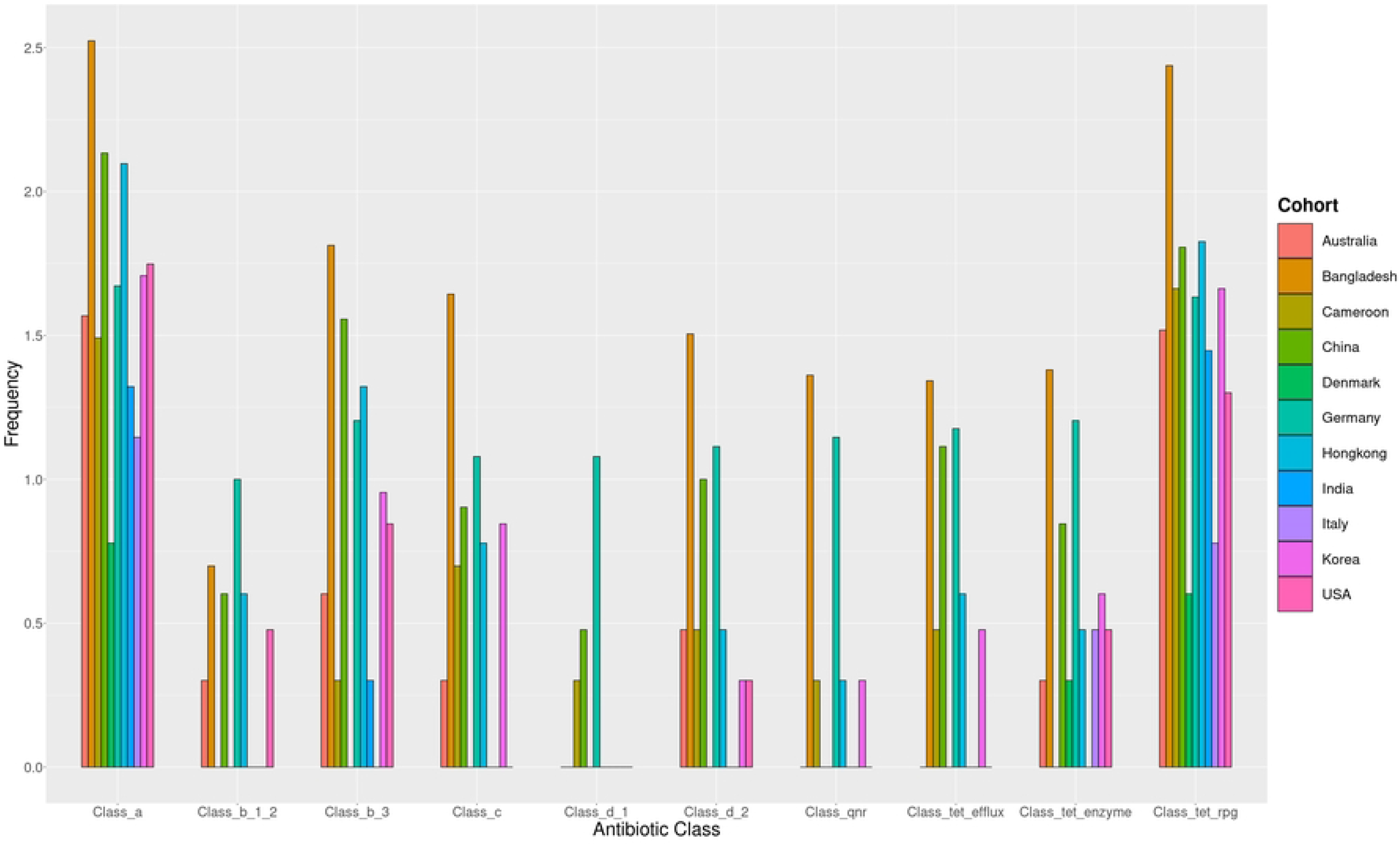
Various antibiotic resistance-causing proteins with their distributions.

**Figure 3:**
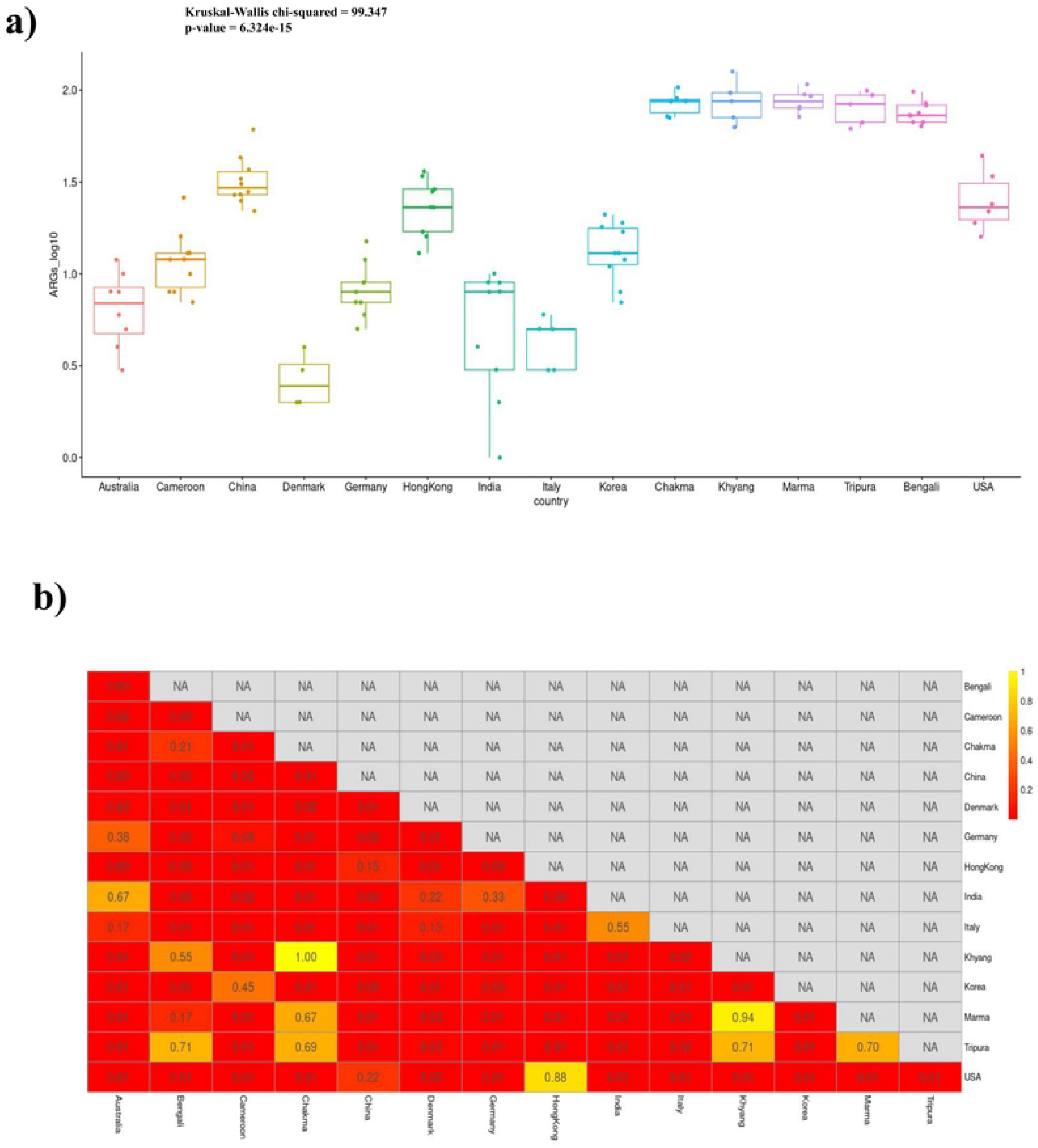
Distribution of antibiotic-resistant genes in each group was visualized using a) Boxplot. Their Kruskal-Wallis test score indicates differences between the cohorts. b) Pairwise Wilcoxon test showing statistically significant p values within the populations. The p values are depicted as a heatmap (score 1 to 0).

### ARGs are more frequent in the mobilome of Bangladeshi samples

The mobile carriers of ARGs are mainly plasmids and viruses (Bacteriophage). Here, the plasmidal and viral DNA sequences were deciphered using several Machine Learning (ML)/ Deep Learning based tools. The detected viral genomes from Bengali, Chakma, Marma, Tripura, and Khyang samples had ARGs on average of 4.4-2.33. The average ARGs of plasmid sequences from Bengali, Tripura, Marma, Khyang, and Chakma were 12-8.33 (**Table 3**).

**Table 3:**
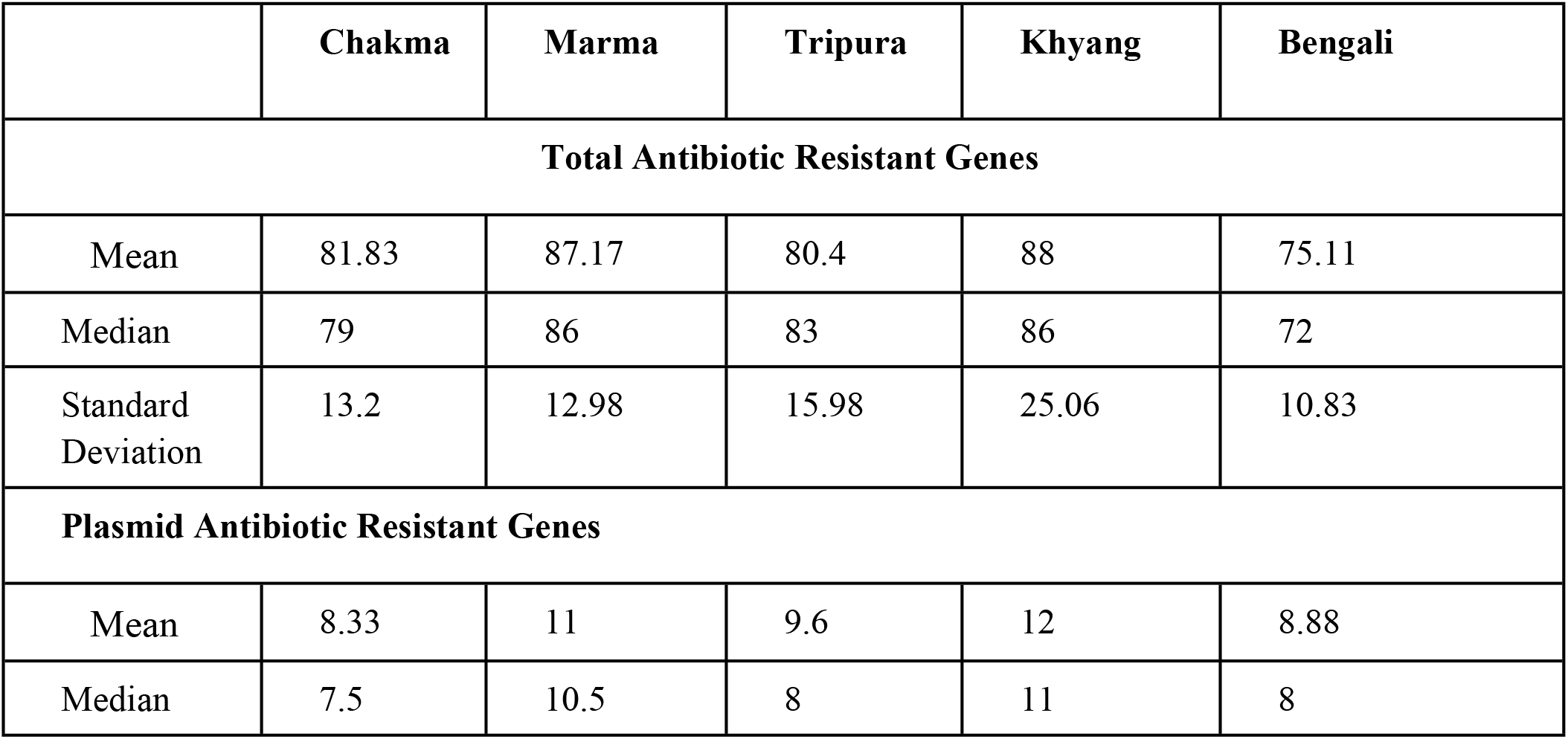
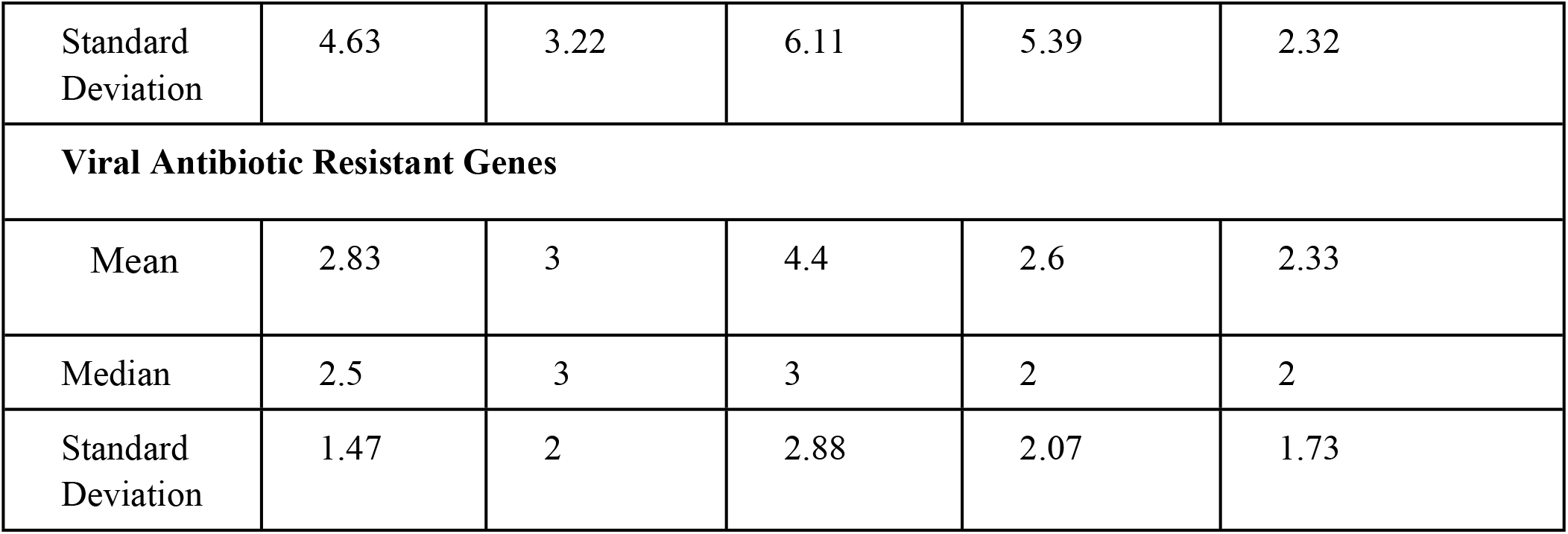
Mean, median, and standard deviation of antibiotic-resistant gene homologs in the total microbiome, plasmidal and viral genomes from Table 1.

In plasmidal sequences catB4, tet(A), tet(W), erm(F), QnrS1, tet(32), tet(O), sul1, vanR-G were highly abundant (**Supplementary Figure 2**). In viral sequences tet(37), tet(44), and tet(O) were more enriched in Bangladeshi virome than in other countries. (**Supplementary Figure 3**). The abundance of the total, plasmids, and viral ARGs was relatively higher in local samples. The number of total ARGs, viral ARGs, and plasmidal ARGs have shown a positive correlation (Spearman’s ρ value: 0.53 & 0.68, p-value < 0.05) (**Figure 4**).

**Figure 4:**
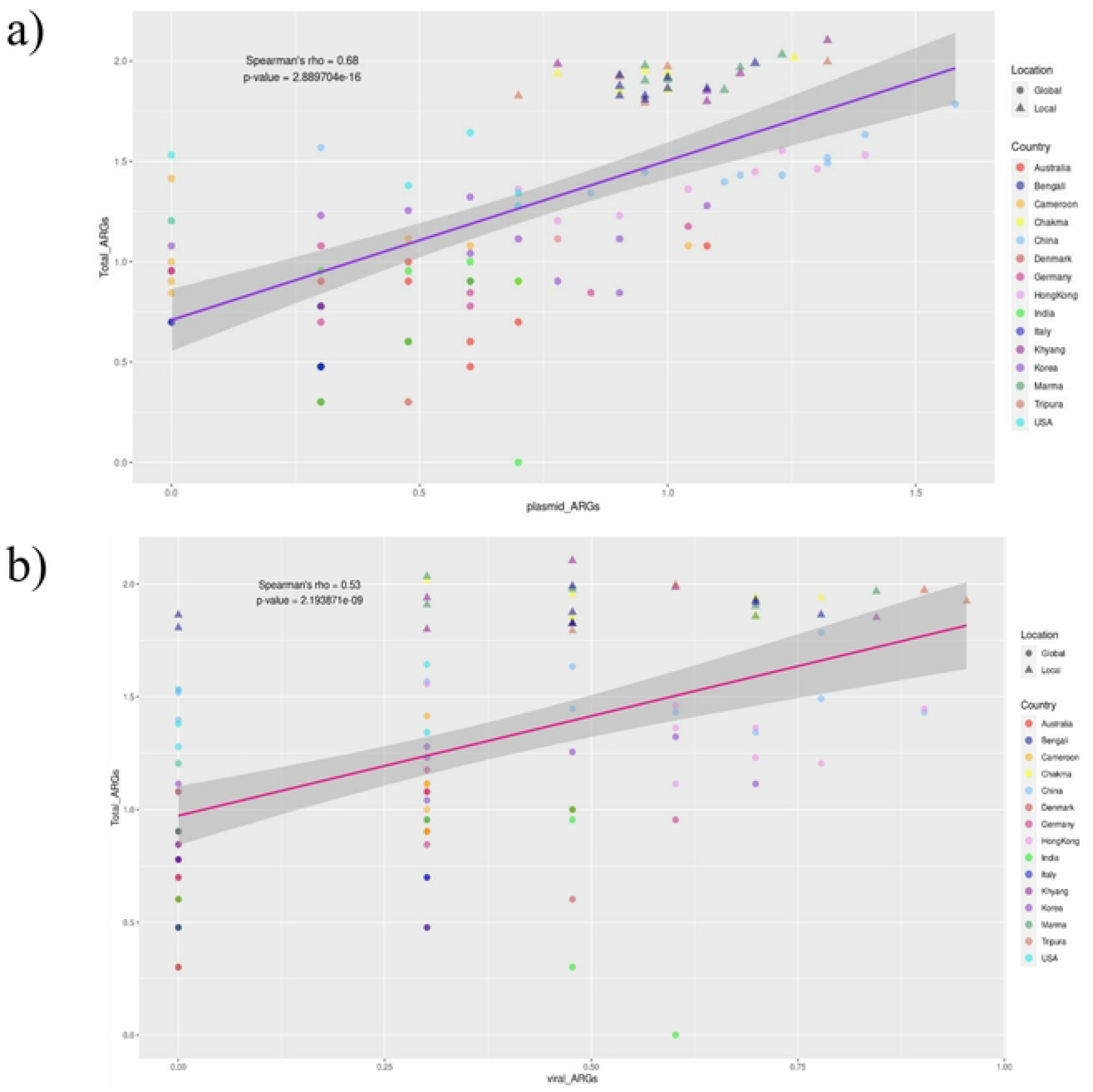
Correlation between total antibiotic-resistant genes with a) Plasmidal antibiotic-resistan genes (Spearman’s rho = 0.68, p-value < 0.05) and b) Viral antibiotic-resistant genes (Spearman’s rho = 0.68, p-value < 0.05).

### Pathway abundance analysis of the selected cohorts

Antibiotic resistance-related pathways were detected using MicrobeAnnotator. The pathways were distributed differently between the cohorts (**Figure 5**). To identify the differentially abundant pathways across all cohorts and each group with Bangladeshi samples, LEFSe was implemented. LEFSe demonstrated differentially abundant pathways in Chakma and Bengali groups. Chakma and Bengali populations have a relatively higher abundance of NorB and MexPQ-OpmE pathways respectively **(Figure 6a)**. However, when all the samples from Bangladesh were joined together and compared with each group, MexPQ-OpmE showed higher abundance in each group except China (**Figure 6b**).

**Figure 5:**
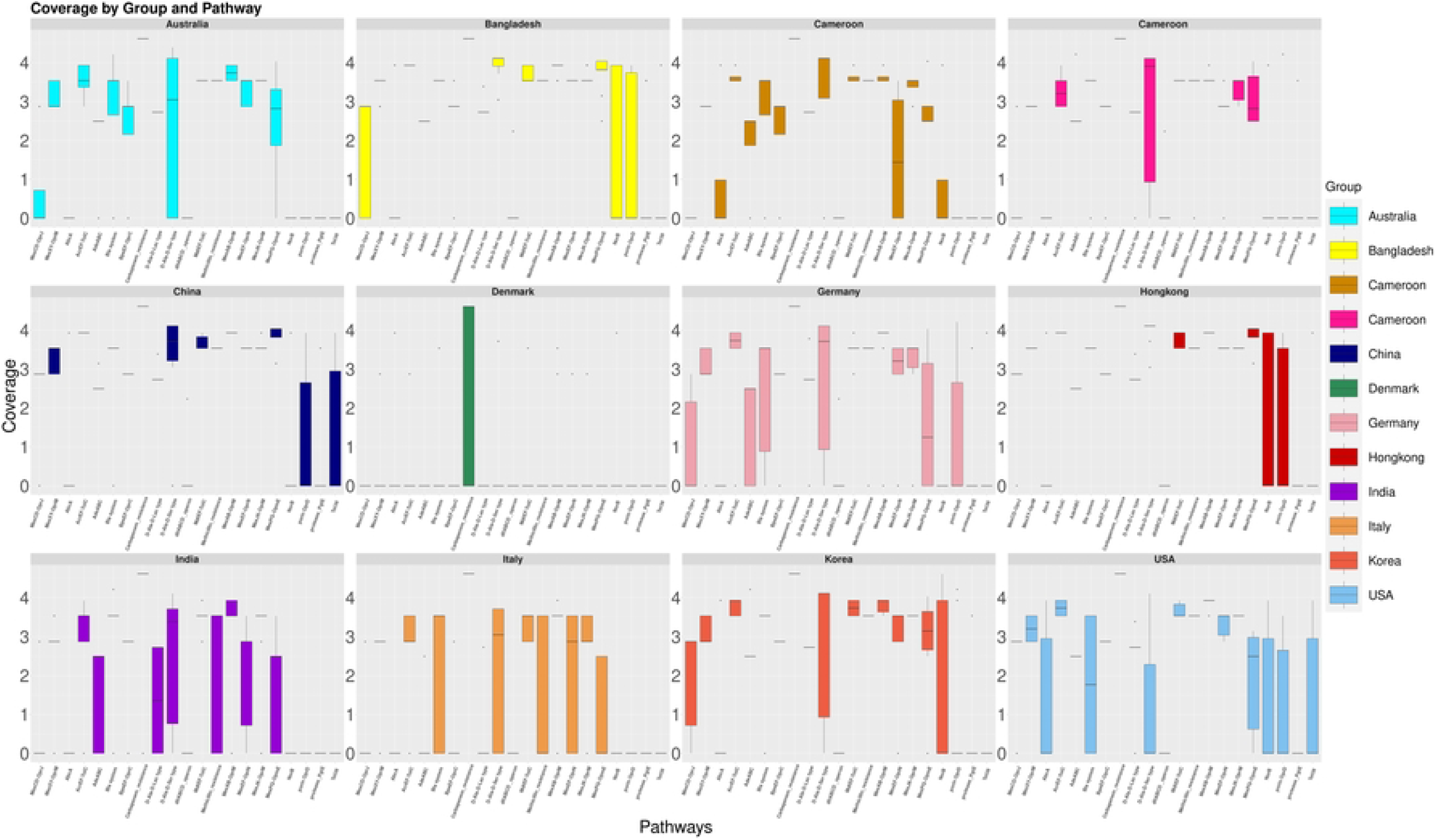
Drug resistance pathway distributions among the groups. Each facet grid represents one group and the distributions of different resistant pathways in those groups.

**Figure 6:**
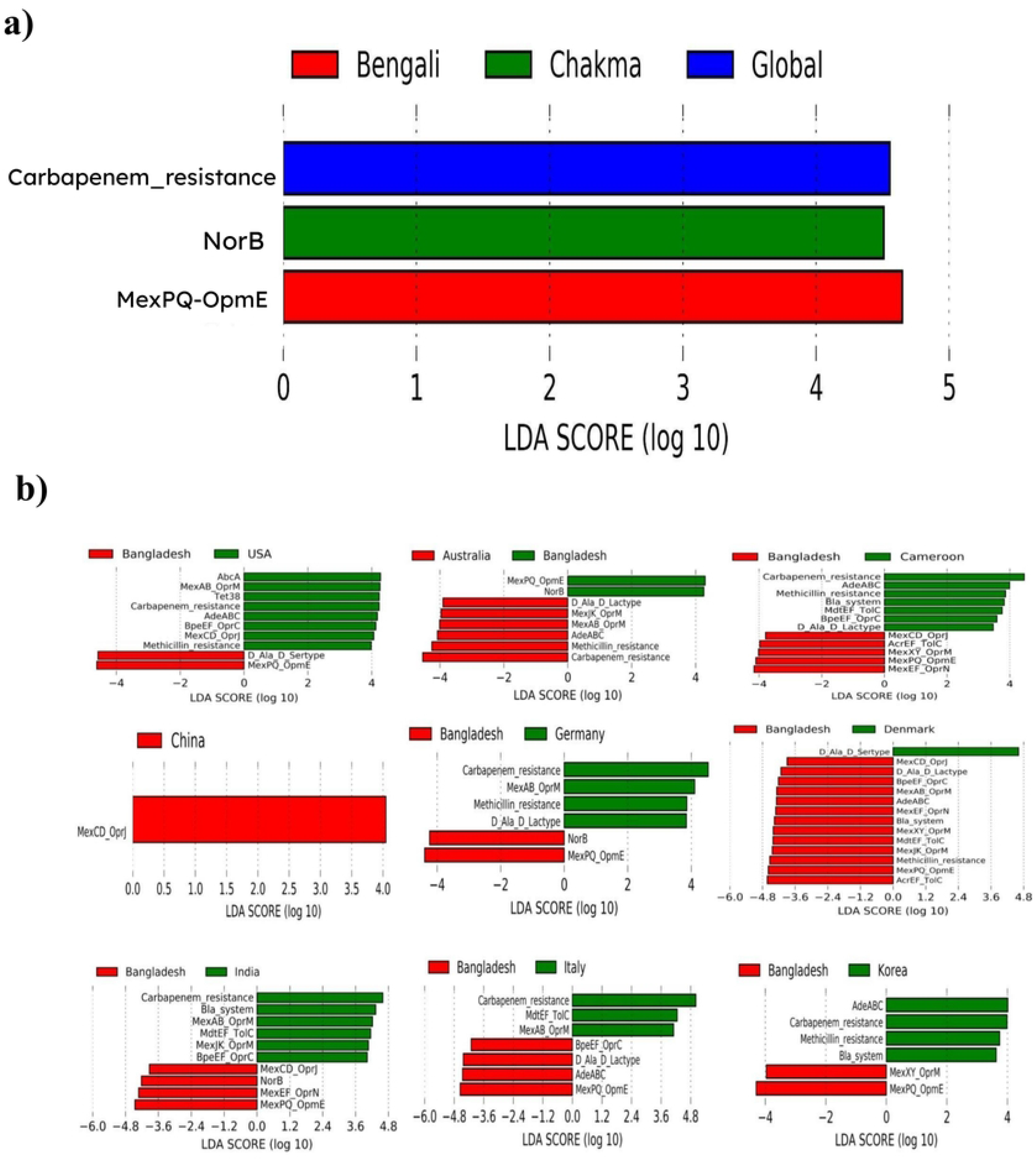
In Linear Discriminant Analysis (LDA) of the pathways from a) all the samples. b) Differentially abundant pathways were also detected for each group with Bangladesh.

## Discussion

Management of Antibiotic pollution in Low and middle-income countries (LMICs) is challenging due to low resources, poor management, and improper infrastructure of sewage systems. In Bangladesh, the prevalence of antibiotic-resistant microorganisms is increasing at an alarming rate (4). In Bangladesh, where industrial development is going in an enormous force, the Bengali and indigenous people are getting used to modern lifestyles for several decades. Since different antibiotics are randomly given over the counter (OTC) in Bangladesh, resistance is booming in this place (29). Moreover, the usage of antibiotics in poultry and animal farms, improper hygiene, and industrial and untreated hospital wastewater are fueling this problem. Such incidents will facilitate the generation of multidrug-resistant pathogens which will be hard to manage (30,31).

Antibiotic resistance and the availability of antimicrobial agents are natural phenomena that have existed since ancient times. Through the episodes of evolution and natural selection, microbes produce antibiotics and bacteria have developed antibiotic resistance to survive against these antimicrobials (32). However, in nature, microbes produce antibiotics at the minute level and antibiotic resistance does not develop rapidly. Enhanced elevation of antibiotics from artificial sources such as medical wastewater, human/ animal manure, and urine can quickly increase the abundance of antibiotic resistance in the environment. Under higher concentrations of antibiotics, a selection pressure will promote turning an ARG-like homolog into a fully functional ARG (32,33). Hence, reservoirs of these ARGs (or resistomes) disseminate ARGs or ARG homologs in the environment and pose a massive threat to public health (33). The human gut microbiome is a vast reservoir of antibiotic resistome (34). Analyzing the human gut resistome is critical to decrypt the spreading of ARGs and their types or prevalence (35). ARGs in non-pathogenic microorganisms are not a threat and sometimes protective for the microbiome against antibiotics, however, these genes can be obtained by pathogenic bacteria through mechanisms such as DNA conjugation, transformation, and transduction (33,36). These processes transmit viral (bacteriophage) elements and mobilomes like plasmids. Therefore, the elevated presence of ARGs in plasmids and viruses indicates more chances of ARG spreading.

In this study, we analyzed the gut resistome of 31 samples from five Bangladeshi cohorts and compared them with different studies. First resistant genes and proteins were detected for the total microbiome. Afterward, the plasmidome and virome were separated. Using ABricate, the resistant genes were identified in these mobilomes. The total abundance of ARGs was higher in local samples. ARGs were also higher in viral and plasmidal elements. The heatmap analysis showed that tet(W), tet(37), catB4, tet(32), VanR_G, erm(X), and many other ARGs were highly abundant in the Bangladeshi population. These types of diversity and elevated abundances create a higher possibility of multidrug-resistant pathogens. However, for other samples (where antibiotics are not available in OTC), these diverse ARGs were less frequent. Log abundance was significantly higher in the local samples and their difference was not statistically significant between them. However, they were different from other countries. Hence, to reduce the ARG abundance proper regulation of antibiotic usage is required..

In plasmidal DNA catB4, tet(A), tet(W), erm(F), QnrS1, tet(32), tet(O), sul1, vanR-G were more frequent in the Bangladeshi cohorts. The same type of pattern was also observed in viral sequences where tet(37), tet(44), and tet(O) were highly enriched in the samples from Bangladesh. The presence of these types of various genes in the mobilome increases the probability of transduction, transformation, and conjugation into pathogenic bacteria. Elevated mobile elements with resistant genes will enhance the generation of drug-resistant pathogens. This pattern was observed in **Figure 4** where we have seen a positive correlation between total resistome and resistant genes in the mobilome. fARGene detected resistance-causing proteins in the whole microbiome and found the elevated frequency of tetracycline efflux pump and tetracycline-resistant related enzymes in Bangladeshi groups. These elevations indicate that tetracycline-resistant genes are spreading at a higher rate in the environment. From the previous studies in Bangladesh, researchers and clinicians have observed that the median tetracycline resistance of *Eschericia coli* and *Staphylococcus aureus* are 65 and 43.5 respectively. For Doxycycline, the value is 61.1 in *E. coli* (37). Hence, these types of broad-spectrum antibiotics might not work in the near future. The fARGene analysis also showed the prevalence of Class b_1_2 beta-lactamase was relatively lower in Bangladeshi samples and Class_d_1 was absent in Bangladeshi samples. Interestingly, Quinolone-resistant genes (Class_qnr) were very high in local samples. Class_qnr was only present in 5 out of 11 groups. Fluoroquinolone such as ciprofloxacin can be found in sewage sludge at milligram/ kilogram concentrations but ciprofloxacin-sensitive strains are very uncommon in sewage (38). Moreover, the administration of Ciprofloxacin did not enrich quinolone resistance (qnr) genes in the gut microbiome (39). However, the presence of qnr type genes in locales most plausibly indicates prolonged exposure to quinolone antibiotics. This exposure might come from the environment, direct consumption of antibiotics, or consumption of foods that contain antibiotics (e.g., Broiler or Layer Chicken) (40). The LEfSe analysis showed a higher abundance of the MexPQ-OpmE pathway in Bangladeshi samples than in other groups. The mexPQ-OpmE pathway may provide fluoroquinolone resistance in the Bangladeshi gut microbiome (41).

Our study suggests controlling the exposure to antibiotic pollution using different regulations from the government and proper authority. Antibiotics in OTC must be halted in Bangladesh to reduce the antibiotic-based selection pressure in the gut microbiome. Moreover, the industrial use of antibiotics should be governed by the authorities properly. We hope our study will elevate consciousness among the people of Bangladesh and other South Asian LMICs.

## Conclusion

Antibiotic-resistant genes are more prevalent in the gut microbiota of Bangladeshi people than in other groups/ countries. This higher prevalence was observed in both urban, suburban, and rural areas. It was detected within different indigenous and Bengali cohorts. The higher ARG frequency in gut mobilome is increasing the possibility of generating multi-drug resistant pathogens using transduction, transformation, and conjugation. Therefore, proper regulation is mandatory in the future to improve this situation.

## Acknowledgments

This study was funded under the “Establishment of National Gene Bank Project”, National Institute of Biotechnology (NIB), Ministry of Science and Technology, Government of the People’s Republic of Bangladesh. We would like to thank Md. Amjad Hossain, Ritaren Chakma, Tina Tripura, Barshan Chakma, and Suborna Dash for their assistance with sample and metadata collection.

## Figure Legends

**Supplementary Figure 1:** Heat map analysis of the ARG abundances in different samples.

**Supplementary Figure 2:** Antibiotic Resistance Gene (ARG) homolog abundances in the gut plasmidome

**Supplementary Figure 3:** Antibiotic Resistance Gene (ARG) homolog abundances in the gut virome

